# Regulation of Aldosterone Secretion by Substance P and the NK1 Receptor in Aldosterone-Producing Adenomas

**DOI:** 10.1101/2025.05.28.656733

**Authors:** Antoine-Guy Lopez, Céline Duparc, Sylvie Renouf, Margot D’Agostino, Kelly De Sousa, Laurence Amar, Guillaume Defortescu, Gilles Manceau, Jean-Christophe Sabourin, Fabio Luiz Fernandes-Rosa, Maria-Christina Zennaro, Tchao Meatchi, Gaël Nicolas, Estelle Louiset, Hervé Lefebvre

**Author notes:** These authors contributed equally to this work (Antoine-Guy Lopez, Céline Duparc). These authors jointly supervised this work (Estelle Louiset, Hervé Lefebvre). **CORRESPONDING AUTHOR** Antoine-Guy Lopez, Univ Rouen Normandie, Inserm, NorDiC UMR 1239, CHU Rouen, Department of Endocrinology, Diabetes and Metabolic Diseases F-76000 Rouen, France.

## Abstract

Aldosterone-producing adenoma (APA) is a major cause of primary aldosteronism (PA), the most frequent form of secondary hypertension. Although somatic mutations in ion channels within APA have been shown to activate Ca^2+^ signaling and drive aldosterone production, the pathophysiology of PA remains partially understood. Substance P (SP), encoded by the *TAC1* gene, is a neuropeptide of the tachykinin family, known for its role in stimulating aldosterone production through activation of the neurokinin 1 receptor (NK1R) in the human adrenal cortex. The aim of our work was to investigate the presence of SP nerve fibers and the NK1 receptor in a large series of APA to assess the potential role of this neuropeptide in the pathophysiology of PA. We analyzed 56 APA tissues using molecular, immunohistochemical, and functional techniques to assess the expression of SP and NK1R and examine the action of SP on aldosterone secretion. SP-positive nerve fibers were detected in 90% of the APA tissues, appearing localized both within and around the adenomas, which also showed strong NK1R expression. Functional studies revealed that SP stimulated aldosterone secretion in 6 of 10 APA cultures. The NK1R antagonist aprepitant inhibited SP-induced aldosterone secretion in 3 of the 4 SP-responsive APA cultures on which the antagonist was tested. Additionally, in perifused APA explants, SP influenced aldosterone pulsatility, resulting in enhanced mineralocorticoid secretion. These findings suggest that the SP-NK1R signaling pathway may contribute to APA pathophysiology and represent a novel potential target for the pharmacological treatment of PA in a subset of patients.

## INTRODUCTION

Primary aldosteronism (PA) is the most frequent form of secondary arterial hypertension with a reported prevalence of 5% in primary care ^1^, increasing to 15-20% in referred populations and patients with treatment-resistant hypertension ^2^. PA is characterized by excessive aldosterone production independent of the renin-angiotensin system (RAS) as evidenced by an elevation of the plasma aldosterone to renin ratio ^3^. Its two major causes are aldosterone-producing adenomas (APAs) and bilateral adrenal hyperplasia (BAH) ^4^. Aldosterone hypersecretion is associated with an increased risk of cardiovascular, renal and cerebrovascular complications ^5^ which resolves after surgical removal of APA or treatment with anti-aldosterone medications ^6,7^.

In 2021, the international HISTALDO consensus further classified unilateral PA into classical and non-classical histopathological types. Classical cases include a solitary APA or a dominant aldosterone-producing nodule, while non-classical cases involve multiple aldosterone-producing nodules, aldosterone-producing micronodules, or diffuse hyperplasia ^8^.

Over the past decade, genomic studies have identified both somatic mutations in genes encoding membrane ion channels and ATPases in up to 90% of APAs and germline mutations in the less frequent familial forms of PA ^9,10^. The affected genes which play a role in aldosterone hypersecretion include *KCNJ5* ^11^, *ATP1A1* ^12^, *ATP2B3* ^12^, *CACNA1D* ^13,14^, *CACNA1H* ^15^, *CLCN2* ^16,17^ and *SLC30A1* ^18^. *In fine*, these molecular alterations result in Ca^2+^ signaling activation in adenoma cells with subsequent increases in both *CYP11B2* (encoding aldosterone synthase) expression and aldosterone production. Conversely, the role of the aldosterone-driver gene mutations in APA development remains unclear, leading to the hypothesis that yet unidentified intracellular pathways may be involved in APA expansion. A possible role of the Wnt/β-catenin pathway has been suggested, based on the observation that β-catenin activation is common in APAs. Although somatic β-catenin mutations, which are usually responsible for activation of the pathway in many tumor types, are less frequently observed in APAs ^19–21^.

Numerous studies have shown that aldosterone production by APAs is not only activated by somatic driver gene mutations but is also regulated by abnormally expressed membrane receptors which confer to adenoma tissues an abnormal sensitivity to various hormones, neuropeptides and conventional neurotransmitters ^22^. These illicit receptors include both overexpressed eutopic receptors which are physiologically present in the zona glomerulosa of the normal adrenal cortex like the serotonin type 4 (5-HT4) receptor and ectopic receptors such as the luteinizing hormone/chorionic gonadotrophin (LH/CG) receptor ^23,24^. Substance P (SP) is a member of the tachykinin family, which also includes neurokinins A and B (NKA and NKB), hemokinin-1, and endokinins. These neuropeptides are involved in pain modulation, emesis, and gonadotropic function and participate in the pathogenesis of menopausal hot flushes ^25–28^. We have recently observed the presence of SP in nerve fibers located in the subcapsular region of the human adrenal gland ^29^. After its release, SP can stimulate aldosterone production through a paracrine mechanism involving the activation of neurokinin type 1 receptor (NK1R), which is expressed by aldosterone-producing cells in the zona glomerulosa. In agreement with this mechanism, administration of the NK1 receptor antagonist aprepitant to healthy volunteers leads to a significant decrease in aldosterone production ^29^. Interestingly, the stimulatory action exerted by SP on aldosterone secretion appears independent and complementary to RAS activation, with SP controlling basal aldosterone production while the RAS triggers aldosterone synthesis in response to upright position ^29^. In addition, integrative omics analyses have shown epigenetic alterations in the NK1R pathway within APA tissues, including hypomethylation of the promotor region of the *TACR1* gene which encodes the NK1 receptor, and concomitant upregulation of *TACR1* mRNA expression ^30^. Based on these findings, we have hypothesized that SP and the NK1 receptor may play a role in the control of aldosterone secretion by APA.

## METHODS

### Patients Samples

We have investigated *in vitro* 56 APAs surgically removed from patients with unilateral PA referred to two French tertiary care centers specializing in adrenal diseases and hypertension, *i.e.* the Departments of Endocrinology and Nephrology of the University Hospital of Rouen, France, and the Hypertension Unit of European Hospital Georges Pompidou, Paris, France. The diagnosis of PA was established according to current guidelines and recommendations ^3,7,31^. Clinical, biological and genetic patient characteristics are summarized in the **Data Supplement** (**Table S1)**. Adrenal samples were obtained at surgery and immediately immersed in culture medium for perifusion and cell culture experiments, frozen at -80°C for RT-PCR analyses, or fixed in formalin for histological studies. Patients included in this study were recruited within the COMETE (COrtico-et MEdullo-surrénale, les Tumeurs Endocrines) - HEGP (Hôpital Européen Georges Pompidou) protocol (authorization CPP 2012-A00508-35). Written informed consent was obtained from all patients for scientific study of adrenal specimens including genetic analyses.

### Real-time RT-PCR

Total RNA was extracted from adrenal tissue using Tri Reagent (Sigma–Aldrich, Saint-Quentin-Fallavier, France) and purified on Nucleospin RNAII (Macherey–Nagel, Hoerdt, France). Control specimens included polyA mRNAs from human adrenal, spinal cord, small intestine and placenta (Clontech, Ozyme, Montigny-le-Bretonneux, France), as well as RNA from LAD2 and Caco2 cell lines (kindly provided by Dr. D Metcalfe, National Institute of Allergy and Infectious Disease, National Institutes of Health, Bethesda, MD; and Dr Moïse Coeffier, Normandie Univ, UNIROUEN, INSERM, Rouen, France; respectively). cDNA synthesis was performed using ImProm-II RT System (SensiFast, Cincinnati, USA). Real-time PCR amplifications were performed with SYBR Green I Master Mix (Applied Biosystem, Courtaboeuf, France) on a QuantStudio 3 System (ThermoFisher, Illkirch, France) using gene-specific primers **(Table S2)**. Each sample was analyzed in duplicates, and cDNA quantification was normalized to *PPIA* (cyclophilin) using the ΔCt Method and standard curves generated from polyA mRNAs or *TACR1* human qPCR Template (HK210362, Origene, Rockville, USA).

### DNA isolation and *KCNJ5* sequencing

Tumor DNA from CYP11B2-positive areas was extracted using QIAamp DNA midi kit (Qiagen, Courtaboeuf Cedex, France) or Maxwell 16 FFPE Plus LEV DNA Kit (Promega). *KCNJ5* was amplified using intron-spanning primers **(Table S3)**. PCR was performed on 100 ng of DNA in a 25 μl reaction volume, containing 0.75 mM MgCl₂, 400 nM of each primer, 200 μM deoxynucleotide triphosphate and 1.25 U Invitrogen Platinum Taq DNA Polymerase (ThermoFisher). The cycling conditions were initial denaturation at 95°C for 5 minutes, followed by 30 cycles at 95°C for 30 seconds, annealing at 60°C, and extension at 72°C for 1 minute with a final extension at 72°C for 7 minutes. Direct sequencing was performed using the ABI Prism Big Dye Terminator® v3.1 Cycle Sequencing Kit (Applied Biosystems, Foster City, CA, USA) on a Hitachi 3500 xL system (Rouen samples) and an ABI Prism 3700 DNA Analyzer (Paris samples).

### Immunohistochemistry

Formalin-fixed, paraffin-embedded adrenal tissue sections were deparaffinized and subjected to antigen retrieval by heating at 95 °C for 20 min in either 10 mM citrate buffer (pH 6) or Tris EDTA (pH 9). Tissue sections were treated with peroxidase blocking reagent (Dako Corporation, Les Ulis, France). Sections were then successively incubated with primary antibodies **(Table S4)** and anti-immunoglobulin antibodies coupled to peroxidase. Immunoreactivities were revealed with diaminobenzidine (Dako Corporation, Carpinteria, CA, USA), and sections were counterstained with hematoxylin. Imaging was performed using an AxioScope 7 microscope (Zeiss) at PRIMACEN, the Cell Imaging Platform of Normandie, University of Rouen Normandie. NK1R expression was semi-quantitatively evaluated by estimating the proportion of stained cells within each sample. A H score was assigned to each tissue based on the percentage of stained cells: undetectable (0, no expression), low (1, <30% positive cells), moderate (2, 30-70% positive cells) and high (3, >70% positive cells).

### Primary cell culture

Ten different aldosteronomas were processed for cell culture experiments. After dissociation with collagenase type 1A and desoxyribonuclease 1 type 4 (Sigma–Aldrich), adrenocortical cells were cultured at a density of 10^6^ cells/ml in culture medium (50% in Dulbecco’s modified Eagle’s medium, DMEM; 50% Ham–F12; ThermoFisher Scientific, Illkirch, France) supplemented with 1% antibiotic–antimycotic solution, 1% insulin–transferrin–selenium solution (ThermoFisher), 10% fetal bovine serum (Sigma–Aldrich) and 10% horse serum (Eurobio-Scientific). To prepare cells for stimulation, the medium was changed 24 hours before treatment, reducing fetal calf serum to 1%. Cells were then incubated in fresh DMEM (basal conditions) or DMEM containing varying concentrations of SP (Sigma-Aldrich) or neurokinin A (Enzo Life Sciences, Lyon, France). SP was administered to cultured cells in the absence or presence of the NK1R antagonist, aprepitant (10^-9^ M; Selleck Chemicals; Houston, USA). A detailed list of the substances tested *in vitro* on aldosterone secretion by APA explants is provided in **Table S5**. Incubation experiments were conducted in quadruplicate at 37 °C in a 5% CO2–95% air atmosphere with 100% relative humidity for 24 h.

### Perifusion experiments

Perifusion experiments were conducted on five different aldosteronomas using a previously described technique ^32^. Tumor explants were dissected into small fragments (1–2 mm^3^), rinsed three times with fresh DMEM and layered into perifusion chambers. Tissue slices were perifused at a constant flow rate of 300 μl/min with DMEM (pH 7.4, 37 °C), continuously gassed with a 95% O_2_-5% CO_2_ mixture. Tissues stabilized for 2 hours prior to the administration of SP (10^-6^ M for 20 min; Sigma-Aldrich). SP was dissolved in gassed DMEM and infused into the perifusion chambers at the same flow rate as DMEM alone by means of a multichannel peristaltic pump. Effluent perifusate fractions were collected every 5 minutes and immediately frozen until aldosterone assay.

### Aldosterone assay

Aldosterone concentrations in culture supernatants and perifusate fractions were quantified using a previously described radioimmunoassay (RIA) procedure with specific antibodies and tritiated hormone (Perkin Elmer, Villebon-sur-Yvette, France) ^33^ and a homogeneous time resolved fluorescence (HTRF) method (Cisbio, Bedford, MA, USA) according to the manufacturer’s protocol. Radioactivity was quantified by using a Tri-Carb 4910TR scintillation counter (Perkin Elmer). Fluorescence was measured by using an Infinite F200 Pro microplate reader (Tecan, Switzerland). Aldosterone concentration was calculated using the sigmoidal standard curve interpolation with Prism 6.0 software (GraphPad Software, Dotmatics; San Diego, CA, USA). For perifusion experiments, aldosterone secretion was normalized to basal level to investigate the action of SP irrespective of spontaneous fluctuations of secretory activity. In this context, basal levels were defined as the average aldosterone secretion measured prior to SP stimulation, serving as a reference for evaluating changes in secretion. Relative changes were expressed as percentages of basal level. RIA and HTRF sensitivities were 80 pg/mL and 25 pg/mL, respectively. Cross-reactivity of aldosterone antibodies with corticosterone, cortisol, testosterone and Δ4-androstenedione were <0.05% for both assay methods.

### Aldosterone Pulse Analysis

Aldosterone secretion pulse analysis was performed as previously described ^34,35^. To evaluate dynamic changes in aldosterone secretion, key parameters were analyzed over two time periods: the basal phase before SP administration (Basal, 0–60 min) and the SP-stimulated phase after SP administration (SP 10^-6^ M, 65–190 min). To account for differences in duration between these periods, the area under the curve (AUC) was normalized per time unit and calculated using Prism (GraphPad Software). Mean aldosterone levels were determined as the average of all measured values. Aldosterone pulse peaks were defined as values exceeding the preceding nadir by ≥10%. Nadir aldosterone levels were calculated as the average of all non-peak values. Pulse frequency was expressed as the number of pulses per hour. Mean pulse interval represented the average time (in minutes) between consecutive peaks. Pulse amplitude was calculated as the difference between each peak and the preceding nadir.

### Statistical analyses

Data were expressed as median ± InterQuartile Range (IQR), minima and maxima values, mean ± SEM, or mean ± SD, as appropriate. All statistical analyses were performed using Prism software (GraphPad Software). The D’Agostino & Pearson test was applied to assess data distribution. Depending on the distribution, statistical significance was assessed by *t* Test, Mann– Whitney *U* test or Kruskal–Wallis test and Dunn’s post-test after one-way ANOVA or two-way ANOVA. Contingency analysis was carried out using the Fisher’s test. Univariate correlations were established using Pearson correlations. P values < 0.05 were considered statistically significant.

## RESULTS

### 1. Expression of tachykinins in APA

We have investigated the expression of the *TAC1, TAC3* and *TAC4* genes which respectively encode SP, neurokinin B and endokinins, by means of quantitative RT-PCR analysis in a series of 42 APA samples. Our results revealed that *TAC1* mRNA levels were significantly higher, compared to those of *TAC3* and *TAC4* mRNAs (P<0.0001), which were either undetectable or barely measurable **(Figure 1A)**. Interestingly, we observed that the expression of the *TAC1* gene was similar among both classical and non-classical APAs (P=0.41), and irrespective of the presence of *KCNJ5* somatic mutations (P=0.33). In addition, statistical analysis of the data shows a moderate positive correlation between APA diameter and *TAC1* mRNA levels, with larger APAs being associated with higher *TAC1* mRNA levels (R=0.41, P<0.01) **(Figure 1B)**. Among the series of 42 APAs, 7 tissues exhibited high *TAC1* expression levels (*TAC1* h) compared to the 35 other samples (*TAC1* l) (P<0.001; **Figure S1).** The 2 subgroups of tissues did not differ in age at surgery, sex distribution, *KCNJ5* mutation status, nodule size, plasma aldosterone levels, or the proportion of classical *versus* non-classical APAs **(Figure S1)**. As depicted in **Figure 1C to G**, approximately 90% of the resected tissues exhibited SP-immunopositive fibers. SP-positive nerve fibers were identified in adenomas and in the adrenocortical tissues adjacent to the adenomas. The distribution pattern of SP-positive nerve fibers was similar in classical and non-classical APAs. Granular SP-immunoreactivity was present in the vicinity of adrenocortical cells, suggesting a potential neurocrine control of endocrine cells. Moreover, a granular cytoplasmic SP staining was detected within a subset of steroidogenic cells. Both SP-positive cells and nerve fibers were observed in CYP11B2-positive areas within adenomas and the adjacent tissues. Additionally, SP-positive fibers and cells were observed within nerve ganglia and along the walls of blood vessels, respectively (**Figure 1H)**.

**Figure 1.**
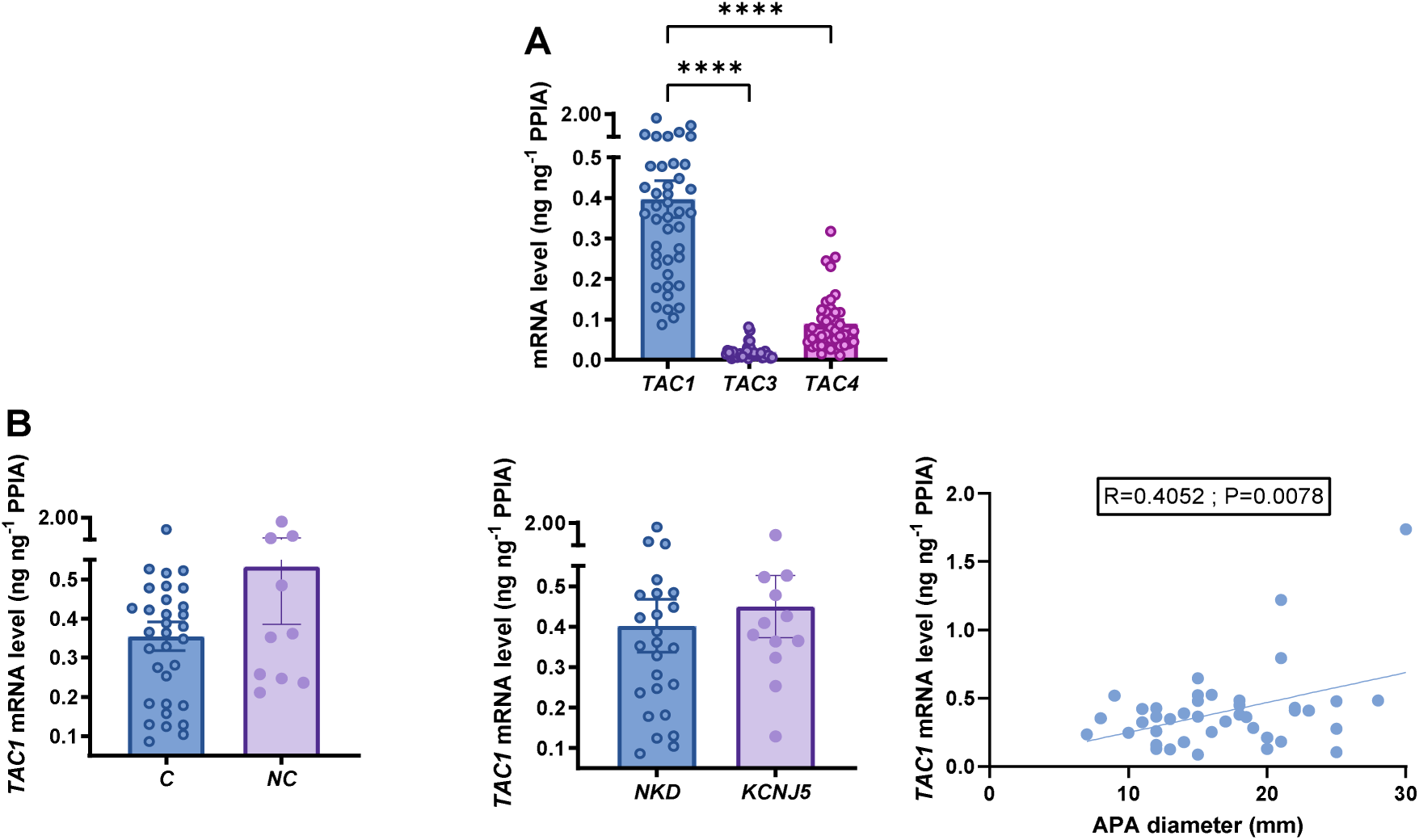
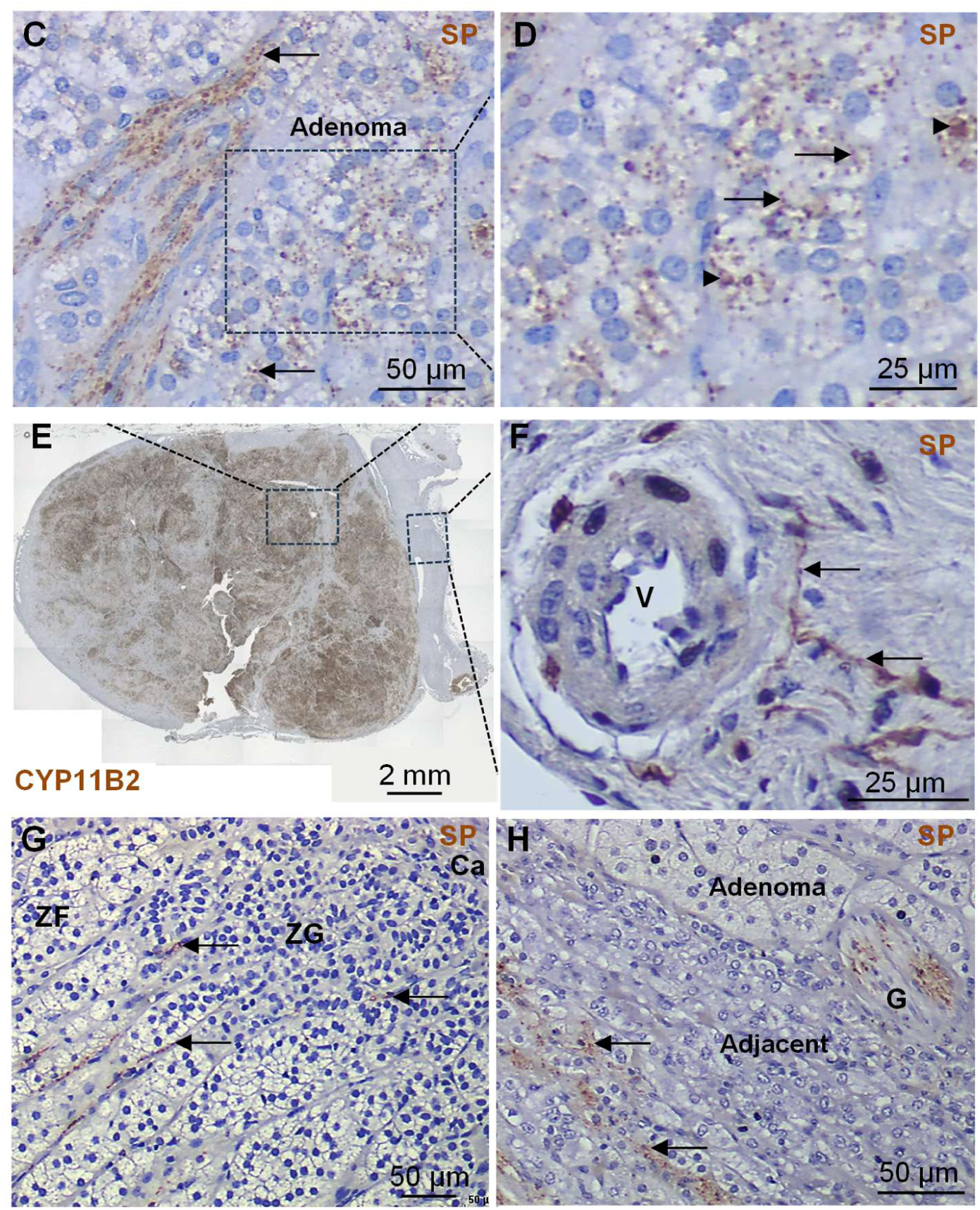
Expression of tachykinins in APA. **A.** Quantitative RT-PCR analysis of *TAC1*, *TAC3* and *TAC4* mRNAs in 42 independent APAs. Each dot indicates one adenoma. Data are presented as mean +/- SEM. ****P< 0.0001. **B.** Comparison of the expression levels of the *TAC1* gene between APA subgroups: C, classical APAs and NC, non-classical APAs (left panel); NKD, no *KCNJ5* mutation detected and *KCNJ5*, *KCNJ5* mutation detected (middle panel), and correlation of the expression levels of the *TAC1* gene with APA diameter (right panel). **C-G.** Immunohistochemical analysis of substance P (SP) in adenoma and adjacent adrenal tissue representative of n = 56 independent APAs. **C-F:** Microphotographs showing a classical APA. **C:** SP-positive longitudinal nerve fibers (arrow) and granulation in the immediate vicinity of steroidogenic cells in an APA region positive for CYP11B2. **D:** High magnification view showing granular SP staining at the periphery (arrow) and within steroidogenic cells (arrowhead). **E:** CYP11B2 immunostaining. The boxed areas highlight the zones of interest shown in C and F panels, exhibiting positive SP staining in both the adenoma (C) and adjacent tissues (F), respectively. **F:** SP-positive fibers (arrows) surrounding a blood vessel (V) in the adjacent adrenal tissue. **G:** SP-positive fibers (arrows) in close contact with steroidogenic cells in the zona glomerulosa (ZG) and zona fasciculata (ZF) in the adrenal tissue adjacent to another classical APA. **H:** SP-staining in the adjacent tissue to the adenoma detected in nerve fibers (arrows) and ganglia (G) in a non-classical APA.

### 2. Expression of tachykinin receptors in APA and correlation with CYP11B2 expression

Next, we investigated the potential effect of SP on aldosterone secretion in APA. The biological actions of tachykinins are known to be mediated by three types of G protein-coupled receptors named NK1, NK2 and NK3 (NK for neurokinin), encoded respectively by the *TACR1*, *TACR2* and *TACR3* genes. Quantitative RT-PCR analysis revealed various levels of *TACR1*, *TACR2*, and *TACR3* mRNA expression. APAs displayed significantly higher expression of *TACR1* mRNA compared to *TACR2* and *TACR3* mRNAs, *TACR3* mRNA levels being nearly undetectable (P<0.05 and P<0.0001, respectively, compared to *TACR1* mRNA levels). In addition, *TACR1* gene expression was significantly higher in non-classical APAs (NC) compared to classical APAs (C) (P< 0.05). *TACR1* mRNA expression levels were similar in *KCNJ5*-mutated and non-mutated adenomas (P=0.79). Within the NC subgroup, *TACR1* mRNA expression levels did not differ significantly between cases with two or more CYP11B2-positive nodules (P=0.07) (**Figure 2A & 2B)**. In order to evaluate potential different impacts of TACR1 and TACR2 on the pathophysiology of APA, we have examined the characteristics of different APA subgroups based on their receptor expression profiles. First, we did not observe any significant correlation between *TACR1* and *TACR2* mRNA levels in both C and NC APAs (**Figure S2**). We have also defined 3 APA subsets according to the *TACR1*/*TACR2* mRNA ratio: *TACR1* predominant expression (*TACR1* p) ratio > 2, *TACR2* predominant expression (*TACR2* p) ratio < 0.5, balanced expression (*TACR1*∼*2*) ratio between 0.5 and 2, respectively (P<0.001; Kruskal-Wallis test) (**Figure S3).** The *TACR1* p subset was exclusively composed of *KCNJ5* non-mutated APAs. In contrast, the 3 subgroups of tissues did not differ in age at surgery, sex distribution, nodule size, plasma aldosterone levels, or the proportion of classical *versus* non-classical APAs. The expression and distribution of NK1R in adrenal tissues were examined using immunohistochemistry in 56 APAs. Heterogeneous NK1R staining was detected in either the adenoma or the adjacent adrenal cortex in 90% of the cases **(Figure 2C to F)**. Out of the 56 APAs, 29 showed high NK1R staining, 9 had moderate staining, and 12 exhibited low staining in the adenoma, according to the H score based on the percentage of stained cells (**Figure S4)**. In the adrenocortical tissues adjacent to the adenomas, NK1R-positive staining was observed in the zona glomerulosa (ZG) and nerve ganglia, as well as in the membranes and cytoplasm of the adrenocortical cells. Additionally, adjacent adrenal micronodules exhibited NK1R immunoreactivity in 27 of the 56 APAs. Interestingly, the comparison between the respective distributions of NK1 and CYP11B2 immunoreactivities within the tissues revealed that both proteins were detected in the same regions in 33 out of 56 cases. Among these adenomas, 22 were classified as classical APAs and 11 as NC APAs, as shown in **Figure 3 & S5.** However, we were unable to clearly define the APA subgroup showing coexpression of NK1R and CYP11B2, since it was not associated with any specific genotype, histological form, or distinct clinical profile.

**Figure 2.**
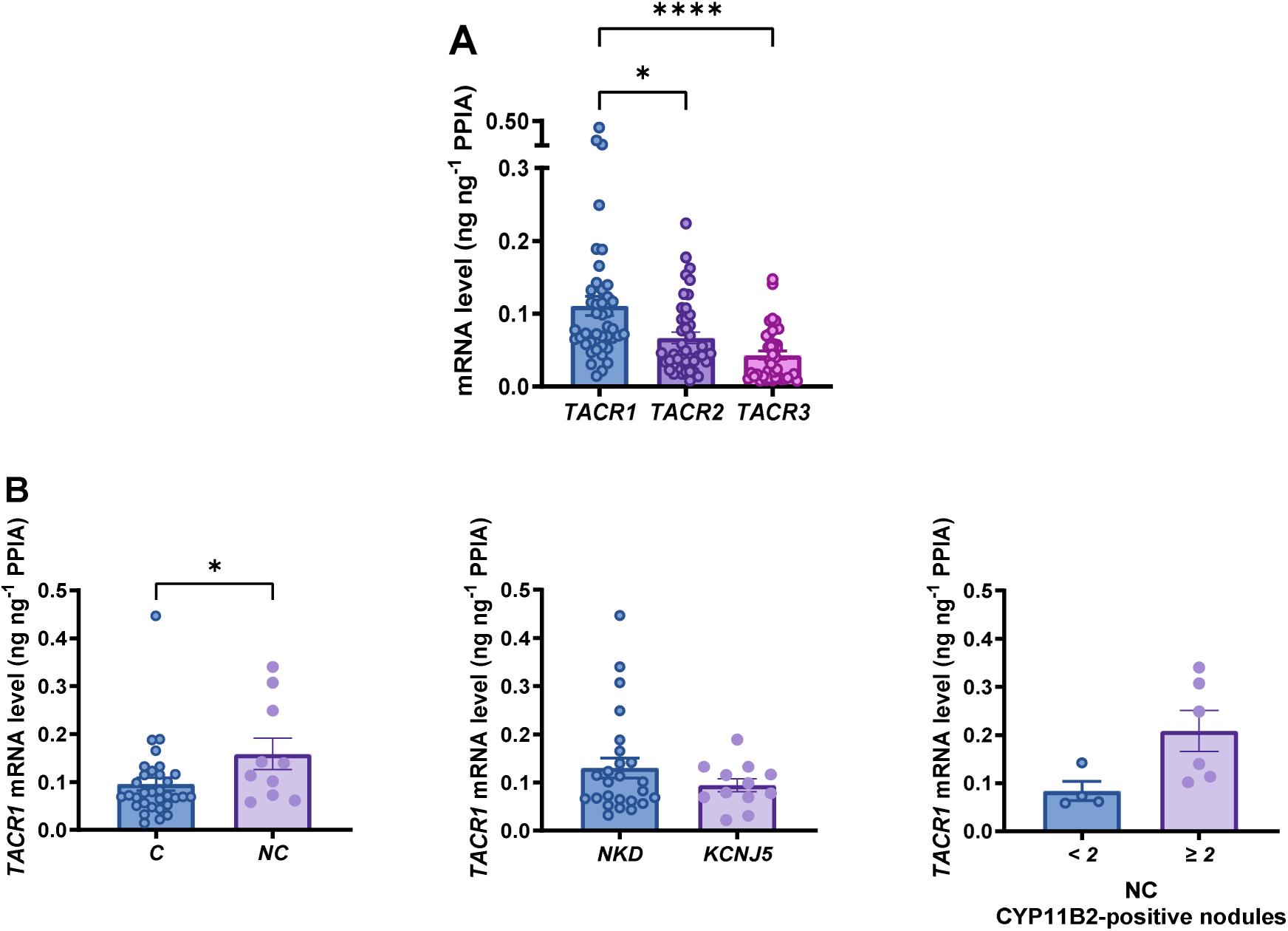
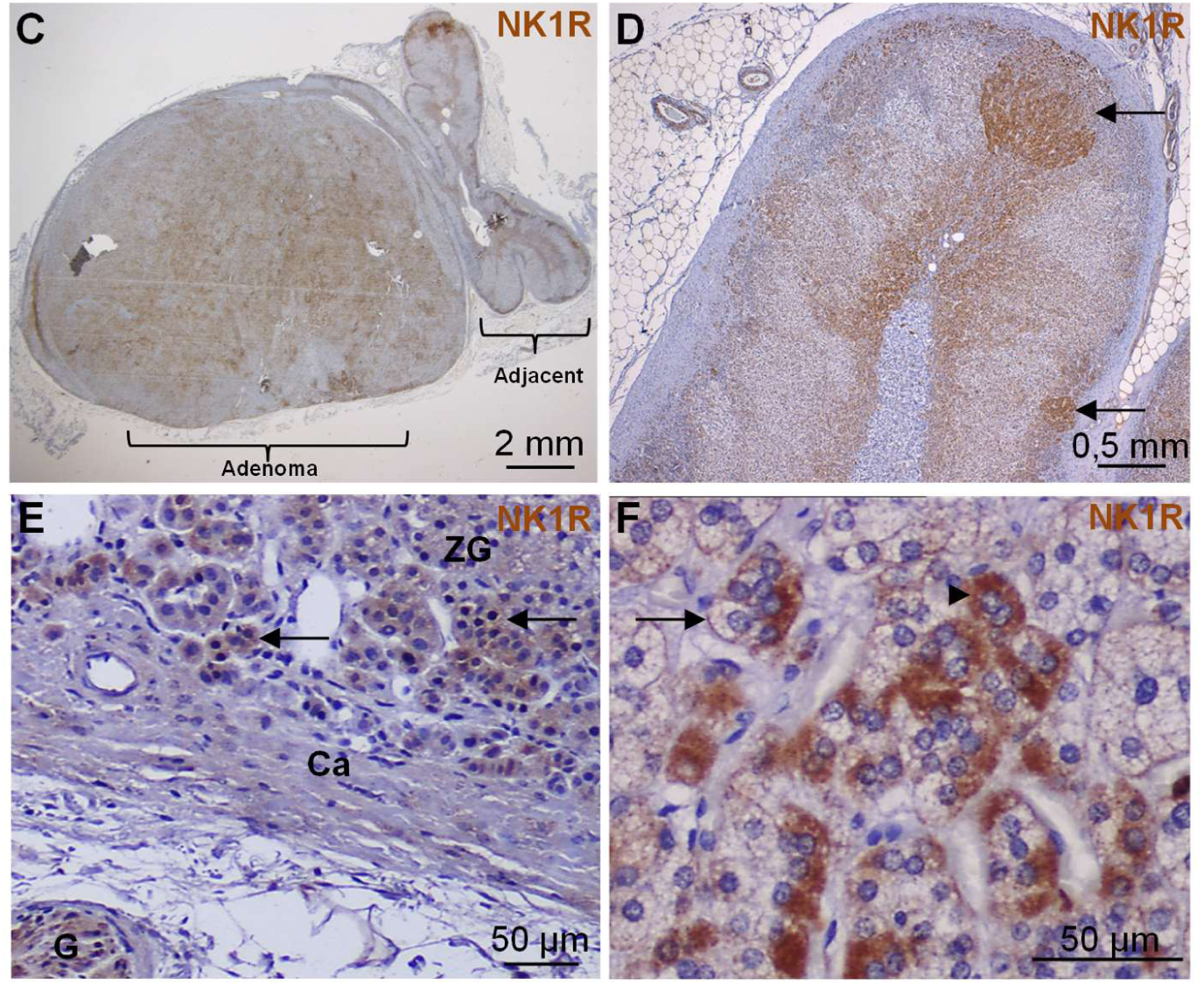
Expression of tachykinin receptors in APA. **A.** Quantitative RT-PCR analysis of *TACR1*, *TACR2* and *TACR3* mRNAs in 42 independent APAs. Each dot indicates one adenoma. Data are presented as mean +/- SEM. *P< 0.05; ****P<0.0001. **B.** Comparison of the expression levels of the *TACR1* gene between different APA subgroups. C, classical APAs; NC, non-classical APAs (left panel); NKD, no *KCNJ5* mutation detected and *KCNJ5*, *KCNJ5* mutation detected (middle panel); CYP11B2-positive nodules in non-classical APAs (< 2 nodules vs ≥ 2 nodules, right panel). **C-F.** Distribution of NK1R immunostaining in APA and adjacent adrenal tissue representative of n=56 independent APAs. **C, E, F:** NK1R staining in a classical APA and its adjacent adrenal cortex. **C:** Overall NK1R staining pattern in an APA and the adjacent adrenal cortex. **D:** Closer view of the NK1R staining in a different classical APA, highlighting areas of staining present in the zona glomerulosa (ZG) or organized in micronodules within the adrenal tissue adjacent to the adenoma (arrows). **E:** Close-up view of NK1R-positive staining in the ZG and nerve ganglia (G) in the adrenal cortex adjacent to the adenoma (Ca, capsule). **F:** High magnification view illustrating the distribution of NK1R staining in a group of adenoma cells. Arrow and arrowhead show membrane and cytoplasmic NK1R staining, respectively.

**Figure 3.**
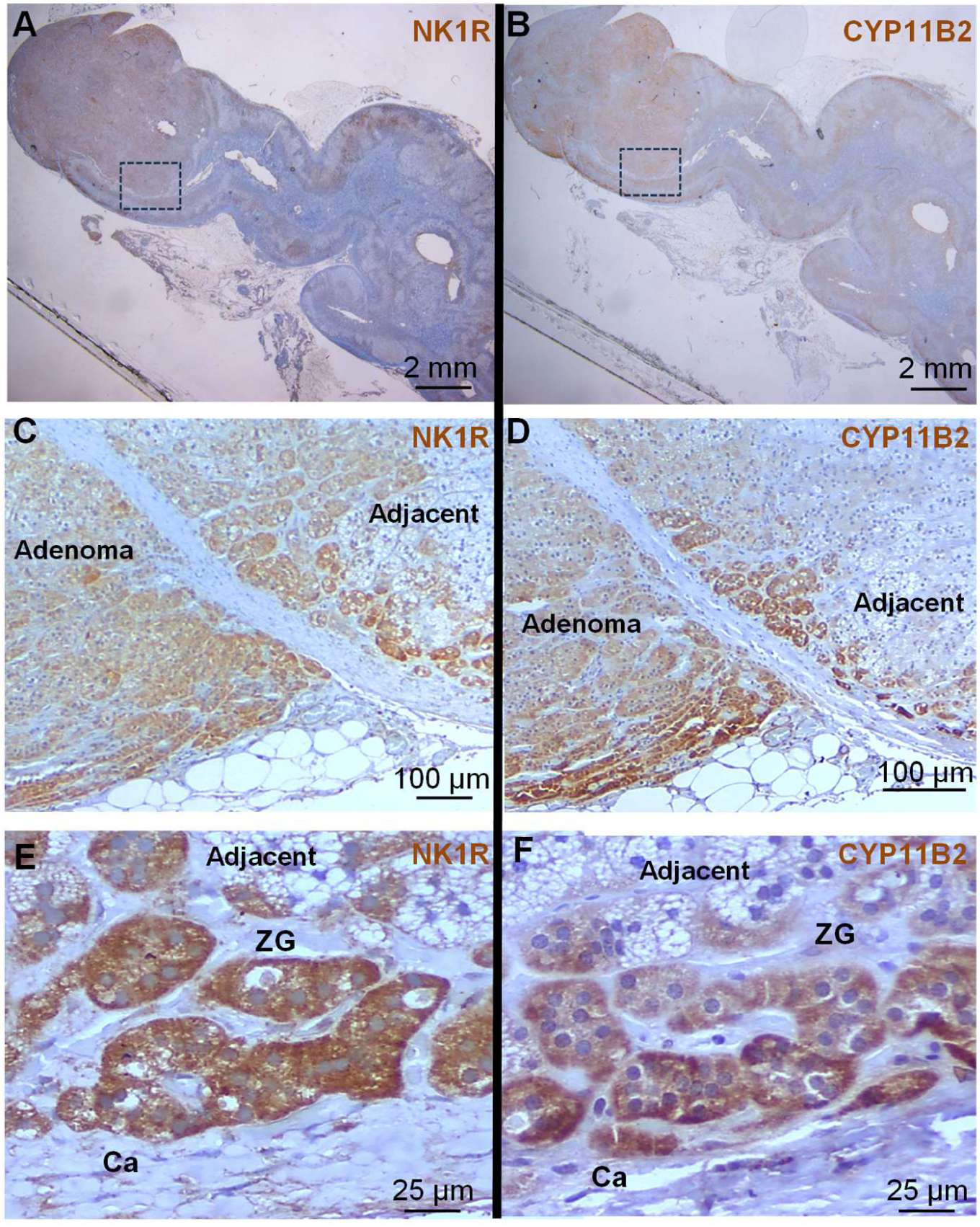
Comparison of NK1R and CYP11B2 expressions in APA by immunohistochemistry. **A to F.** Comparison of NK1R and CYP11B2 immunostainings in the adenoma and adjacent adrenal tissue, illustrated by microphotographs representative of n = 33 independent APAs. All panels depict a classical APA. **A-B**: Low magnification views of NK1R (A) and CYP11B2 (B) immunostainings in the same APA tissue. **C-D**: Closer view of NK1R (C) and CYP11B2 (D) staining patterns in adenoma and adjacent adrenal cortex. **E-F**: Higher magnification views showing NK1R (E) and CYP11B2 (F) staining in the zona glomerulosa (ZG) and adrenal capsule (Ca) in the adjacent tissue.

### 3. Effect of tachykinins on aldosterone production from APA tissues

We investigated the effects of SP, which primarily binds to and activates the NK1 receptor, and NKA, which preferentially binds to and activates the NK2 receptor, on aldosterone production by APA cells in primary culture and perifused APA explants (**Figure 4**). The impact of SP on aldosterone production was examined in 10 independent APA cell cultures **(Table S5)**. SP (10^-7^ M) stimulated aldosterone production by cultured cells derived from 6 APA (Patients: P1, P4, P5, P6, P8, P10; mean ± SEM: Basal: 100 ± 4.9 %, CI%: 87.4–113.6 *versus* SP, 147.2 ± 13.3%, CI 95%: 97–173; P<0.01) **(Figure 4A)**. Among them, SP dose-dependently stimulated aldosterone secretion in one case (P1; EC50 = 1.71 ± 0.3 nM, Emax 228.9 ± 14.3 % of basal level) **(Figure 4B)**.

**Figure 4:**
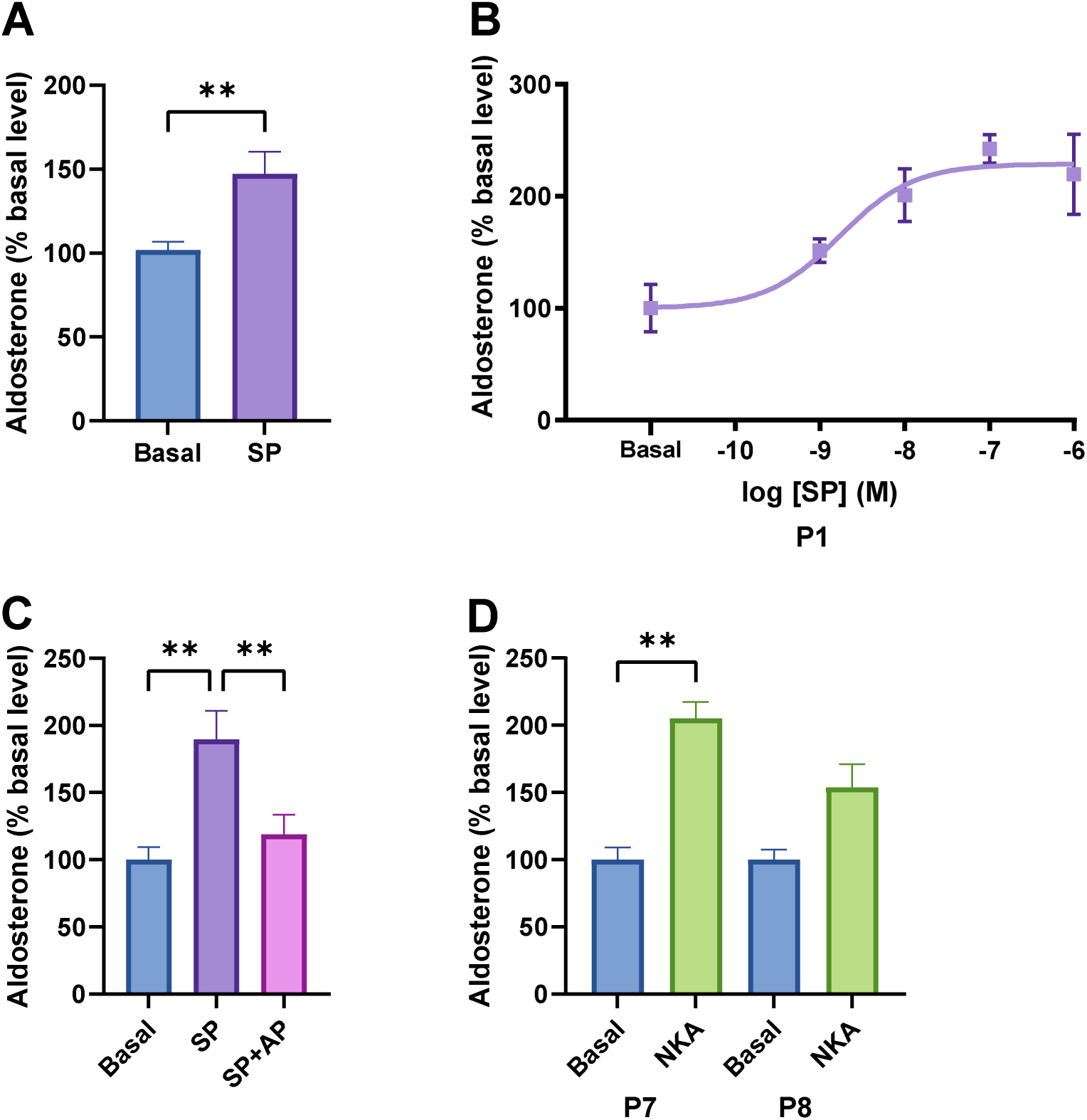
Effect of tachykinins on aldosterone production by APA cells in primary culture. **A.** Effect of SP on aldosterone production by cultured APA cells (n = 10 independent APA cell cultures, each performed in quadruplicate) (**P<0.01). SP (10^-7^ M) significantly increased aldosterone production in 6 APA samples. **B.** Effect of increasing concentrations of SP (10^-10^ M to 10^-6^ M) on aldosterone production in one of the 6 SP-sensitive APAs (from patient 1 [P1]), showing a dose-dependent increase in aldosterone secretion (EC50 = 1.71 ± 0.3 nM, Emax 228.9 ± 14.3 % of basal level). **C.** Effect of SP (10^-7^ M) alone or in the presence of the NK1R antagonist aprepitant (AP; 10^-9^ M) on aldosterone production by cultured APA cells (n = 3 independent cultures). The stimulatory effect of SP was blunted by AP (**P<0.01). **D.** Effect of neurokinin A (NKA; 10^-7^ M) on aldosterone production by cultured APA cells (n = 2 independent cultures from patients 7 [P7] and 8 [P8]). NKA significantly increased aldosterone production in the APA cell culture derived from P7 (**P<0.01), but did not significantly modify aldosterone release in the cell culture derived from P8 (P=0.084). Data represent the mean ± SEM of the values obtained in independent experiments and are expressed as % of basal level. In all culture experiments, aldosterone secretion was normalized to basal levels.

The effect of the NK1R antagonist aprepitant on SP-induced aldosterone secretion was evaluated in 4 SP-responsive APAs. The aldosterone response to SP was inhibited by aprepitant (10^-9^ M) in 3 cases (P1, P4, P5; Basal: 100 ± 9.5 %, CI%: 75.7–142.7 *versus* SP 10^-7^ M, 189.7 ± 21.3 %, CI 95%: 107.5– 249.4, P<0.01; SP (10^-7^ M) + aprepitant (10^-9^ M), 119 ± 14.4 %, CI 95%: 75.2–180.3; P<0.01 *versus* SP alone) **(Figure 4C)**. The effect of NKA was studied in 2 independent APA cell cultures. NKA stimulated aldosterone secretion in one APA, which was unresponsive to SP (P7; Basal: 100 ± 9 %, CI%: 77–120 *versus* NKA (10^-7^ M), 205.2 ± 12.3 %, CI 95%: 180.6–233.7, P<0.05) **(Figure 4D)**. In the APA unresponsive to NKA, SP alone significantly increased aldosterone secretion (P8; Basal: 100 ± 7.7 %, CI%: 82.6–113.6 versus NKA (10^-7^ M), 153.8 ± 17.4 %, CI 95%: 122.9–200.2, P=0.08 and basal *versus* SP (10^-7^ M), 141.1 ± 11.7 %, CI 95%: 119.2.6–173.4, P<0.05). The effect of aprepitant was not studied in this case. Among APAs, the responsiveness to SP was not associated with any specific genotype, histological form, or clinical profile.

The kinetics of the aldosterone response to SP was further investigated using the perifusion technique in tissues derived from 5 different APAs. In all cases, aldosterone secretion spontaneously exhibited a pulsatile pattern **(Figure S4)**. Analysis of the data showed that the mean aldosterone level after SP administration was not different from that observed in basal conditions (Basal = 100 ± 5 vs. SP = 128 ± 16.8 % basal level; P = 0.148) **(Figure 5A)**. The nadir aldosterone levels remained similar during the two conditions (Basal = 100 ± 4.9 vs. SP = 112.7 ± 6.9 % basal level; P = 0.173) (**Figure 5B**). There were no statistically significant changes in pulse frequency (Basal = 2.7 ± 0.67 vs. SP = 4 ± 0.58 pulses/h; P = 0.167) **(Figure 5C)**, mean aldosterone pulse interval (Basal = 9.3 ± 2.8 vs. SP = 11.6 ± 1.9 min; P = 0.52) **(Figure 5D)**, or aldosterone pulse amplitude (Basal = 100 ± 31.5 and SP = 219.7 ± 70.2 % basal level; P = 0.158) **(Figure 5E)**. Conversely, the integrated aldosterone response, calculated as the area under the curve (AUC), showed a significant increase after SP administration (Basal = 100 ± 22.8 and SP = 252.8 ± 46.9 % basal level x min; P<0.05) **(Figure 5F)**.

**Figure 5:**
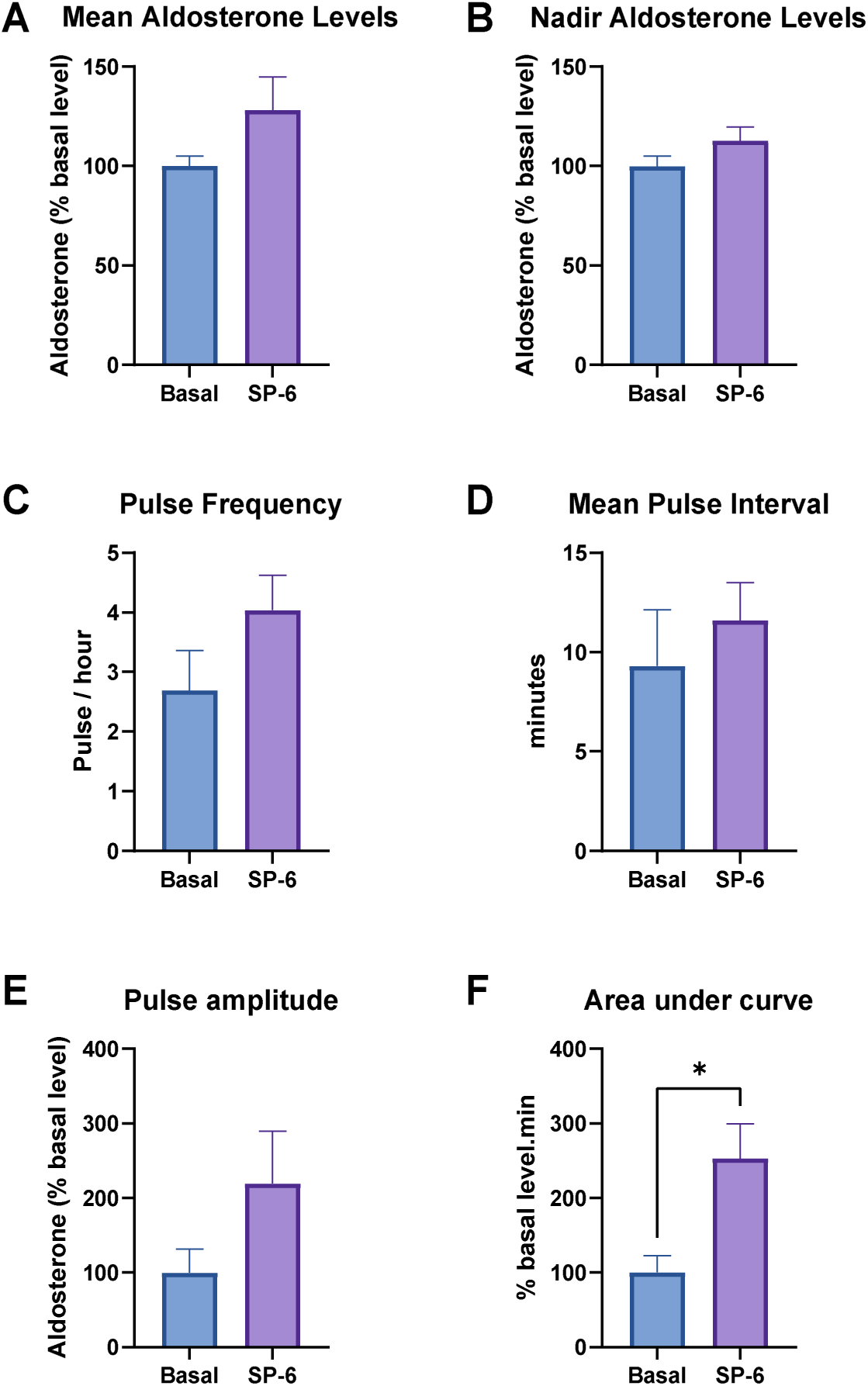
Effect of SP on aldosterone production by perifused APA Tissues. Aldosterone production kinetics in basal state and in response to SP (10^-6^ M, administered for 20 minutes) in 5 independent perifused APA tissues. Perifusion experiments were conducted over a total duration of 250 minutes. **A.** Integrated aldosterone response using the area under the curve (AUC). **B.** Mean aldosterone levels calculated as the average of all values during each condition. **C.** Nadir aldosterone levels, representing the lowest non-peak values in each condition. **D.** Aldosterone pulse frequency expressed as the number of pulses per hour. **E.** Mean aldosterone pulse interval, defined as the average time (in minutes) between consecutive pulse peaks. **F.** Pulse amplitude, calculated as the difference between each peak and its preceding nadir. For each tissue, the data were normalized to basal secretion levels to allow evaluation of the aldosterone response to SP irrespective of the variability of the spontaneous aldosterone production among APAs. Error bars represent the standard error of the mean (SEM). Data are expressed as % basal level.

## DISCUSSION

Our team has recently demonstrated that SP, a neuropeptide localized in subcapsular nerve fibers of the human adrenal gland, stimulates aldosterone production *via* activation of the NK1 receptor, this regulatory mechanism being independent of the RAS system. Based on these findings, we have hypothesized that SP and the NK1R may play a role in the control of aldosterone secretion by APA.

Our results show that *TAC1* mRNA is predominantly expressed in APAs, suggesting that SP may be synthesized by these tumors. In contrast, *TAC4* mRNA is detected at low levels, and *TAC3* mRNA is undetectable, making *TAC1* the primary potential source of SP production ^36^. This pattern of tachykinin gene expression is thus similar to that observed in the normal adrenal gland ^29^. SP-positive fibers were first observed in the normal adrenal cortex by Heym *et al.* but were not precisely characterized ^37,38^. Our previous work showed that intracortical SP-positive fibers are independent of adrenergic and cholinergic fibers, adding further complexity to adrenal innervation. This finding is consistent with the known role of SP as a key neuromediator of the non-adrenergic, non-cholinergic (NANC) system, which is often considered as a third component of the autonomic nervous system although usually operating in synergy with the sympathetic system ^29,39^. We now show the presence of SP-containing nerve fibers in APAs, located both in direct contact with adenoma cells as well as around peripheral blood vessels. These localizations suggest that SP may regulate aldosterone secretion through a dual mechanism, including a direct action on steroidogenic cells and an indirect effect through modulation of adrenal blood flow. Indeed, as SP is a known vasodilator peptide ^40^, its release near adrenal blood vessels may enhance adenoma perfusion. In addition, it is well established that the adrenal blood flow is a crucial determinant of steroidogenesis. As a matter of fact, increased adrenal blood flow, induced by vasodilatory agents like histamine, has been shown to enhance corticosteroid production in rats ^41^. This stimulatory effect on aldosterone production could be attributed to an increased availability of substrates essential for corticosteroidogenesis, such as cholesterol ^42^. Alternatively, Vinson *et al*. have proposed that enhanced blood flow could activate the production of endothelin, a potent stimulator of aldosterone secretion, by adrenal capillary endothelial cells ^43^. Our immunohistochemical studies also showed the presence of detectable amounts of SP in the cytoplasm of APA adrenocortical cells. The granular appearance of the staining suggests that the peptide is stored in secretory granules in agreement with the well-established neuroendocrine differentiation of APA tissues ^44–46^. Collectively, these results provided a histological basis for an autocrine and neurocrine/paracrine regulation of aldosterone secretion by SP in APAs.

We found that *TACR1* mRNA is highly expressed in APAs, while *TACR2* expression is weaker but still significant and *TACR3* mRNAs appear undetectable in the great majority of tissues. This contrasts with the normal adrenal gland, where *TACR2* mRNA levels are barely detectable ^29^ indicating that NK2R may be considered as an aberrantly expressed receptor in APAs ^47^. Our findings are supported by previous transcriptomic analyses that revealed overexpression of *TACR1* and *TACR2* in 48 APAs compared to normal adrenal tissues ^48^. Consistently, Itcho *et al.* reported *TACR1* mRNA overexpression in APAs compared with non-functioning adrenal adenomas ^30^. Our immunohistochemical analysis of NK1R distribution corroborated the mRNA expression data by showing diffuse and heterogeneous NK1R labeling in both adenomas and adjacent adrenal cortex in most tissues. The close proximity of NK1R-expressing adrenocortical cells to SP-containing nerve fibers, combined with the widespread distribution of NK1R within tumor tissue, suggests that SP may exert direct control over on aldosterone production via NK1R. We have thus investigated the possible co-expression of NK1R and CYP11B2. Immunohistochemical data show that the two proteins localize in the same groups of adrenal cells in a subset of APAs which represents the majority of the tumors studied. However, this subgroup of APAs does not seem to be associated with a specific genotype, distinct histological form, or any particular clinical characteristics.

The next step was to investigate the effect of SP on aldosterone secretion using both primary cultures of APA-derived cells and perifusion of APA fragments. SP increased aldosterone production in 6 out of 10 tumors. This variability in the secretory response probably results from the heterogeneous expression of the NK1R within APAs. As circulating SP levels in human plasma generally do not exceed 10^-11^ M ^49^, a concentration insufficient to stimulate aldosterone secretion *in vitro*, SP regulation of aldosterone production by APAs likely involves peptide release from adrenal nerve endings or adenomatous cells. Under our experimental conditions, the maximum efficiency (Emax) of SP on aldosterone secretion reaches around +50% of the basal level, a lower response compared to the 70% increase observed in normal adrenal tissue ^29^. This reduced responsiveness of APA tissues to SP might be due to receptor desensitization ^50^, or partial saturation by endogenous SP in APA cells ^28^. The EC50 of SP to stimulate aldosterone secretion was consistent with the known affinity of SP for NK1R, supporting the involvement of this receptor in SP-induced aldosterone production in APA cells ^51^. Accordingly, the NK1R antagonist aprepitant inhibited SP-induced aldosterone secretion in the majority of the APAs studied, confirming NK1R involvement in a subpopulation of APAs, consistent with its role in normal adrenocortical physiology ^29^. Additionally, NKA, which binds to NK2R, significantly stimulated aldosterone secretion in one APA cell culture that was unresponsive to SP. This suggests that NKA, produced by the *TAC1* gene alongside with SP, may regulate aldosterone production *via* the NK2R in some APAs. In addition, it cannot be excluded that SP may also activate the NK2R expressed by some APAs. In fact, although it is known that SP binds the NK2R only at high concentrations (10^-6^ M or above), the combined production of the peptide by nerve fibers and some adenoma cells may generate local concentrations capable of stimulating the NK2R signaling pathway ^52^.

Our findings reveal that aldosterone secretion by perifused APA tissues exhibits spontaneous pulsatility, occurring thus independently of any external stimuli. Interestingly, Siragy *et al.* had demonstrated *in vivo* that plasma aldosterone concentration in APA patients shows regular pulses, including predominant peaks every two hours superimposed with additional micropulses appearing approximately every 30 minutes, a frequence relatively close to that of aldosterone pulses observed in our *in vitro* model (one pulse every 20 minutes) ^53^. It seems therefore likely that this microcyclicity reflects the intrinsic pulsatility of the APA secretory activity, as evidenced by the perifusion technique. The mechanisms underlying the APA intrinsic pulsatility may be multiple, including the expression of clock genes by adrenal cortical cells ^54,55^. Local regulatory loops within the tissue, potentially governed by autocrine or paracrine signaling, may also contribute to this secretory pattern. While the precise regulation of the *in vivo* aldosterone rhythmicity has not been extensively studied, similarities can be drawn with cortisol regulation ^56^. Cortisol rhythmicity is mainly controlled by the pituitary gland and the suprachiasmatic nucleus which plays a central role in the response to stress and light. There is also evidence suggesting that sympathetic innervation of the adrenal cortex contributes to this synchronization, functioning independently of pituitary control to coordinate central and adrenal clocks ^57,58^. Thus, it is plausible that the nervous adrenal command mediated by SP plays a role in the secretory rhythm of APAs. Supporting this hypothesis, our perifusion experiments shows that SP administration influences the pulsatility of aldosterone secretion as evidenced by the increase of the area under the curve which reflects the global SP-induced production of aldosterone. The fact that the different parameters of rhythmicity, *i.e.* mean aldosterone level, nadir, pulse frequency, amplitude, and peak intervals, did not exhibit any significant changes after SP administration probably results from the limited number of perifused APAs together the well-known heterogeneity of the tissues.

In conclusion, our data indicate that tachykinins, especially SP expressed in APAs, are able to stimulate aldosterone production by adenoma tissues through an autocrine and/or neurocrine/paracrine mechanism mainly involving the NK1 receptor.

## PERSPECTIVES

NK1R antagonists like aprepitant which is currently used for the treatment of nausea induced by anticancer chemotherapies, may represent a new pharmacological treatment of APA-related primary aldosteronism in a significant subpopulation of patients. Eligibility of the patients to this new therapeutic approach could be first assessed on the basis of an aprepitant suppression test to verify whether aldosterone secretion can actually be reduced by antagonizing the NK1R.

## ACKOWLEDGMENTS

We are grateful to the French Cortico et Medullo-Surrénale: les Tumeurs Endocrines (COMETE) network and the Tumor BioBank-Biological Resource Centre of Rouen University Hospital directed by Prof. J.-C. Sabourin for providing tissue samples. We also thank Dr C. Gomez-Sanchez (University of Mississippi Medical Center, Jackson, MS) for his generous gift of CYP11B2 antibodies. We acknowledge Dr. D. Metcalfe (National Institute of Allergy and Infectious Diseases, National Institutes of Health, Bethesda, MD) for kindly providing the human mast cell line LAD2, and Dr. M. Coeffier (Univ Rouen Normandie, INSERM U1073, Rouen, France) for the Caco2 cell line. We also thank Dr. N. Sarafan-Vasseur and S. Rousseau (Univ Rouen Normandie, INSERM U1245, Rouen, France) for their support with tissue sequencing; Dr. F. Blanchard (Tumor BioBank, CHU Rouen, France) for her assistance with biobanking logistics; Drs J. Riancho and I. Belmihoub (Hypertension Unit, AP-HP, Hôpital Européen Georges Pompidou, Paris, France) for their valuable assistance in collecting clinical data; and Dr. S. Boulkroun (Université de Paris, Paris Cardiovascular Research Center, INSERM, Paris, France) for her help in preparing adrenal tissue slides. The images were obtained at PRIMACEN (the Cell Imaging Platform of Normandy, University of Rouen, Rouen, France).

## Abbreviations

APA: aldosterone-producing adenoma
NK1R: neurokinin type 1 receptor
NKD: no *KCNJ5* mutation detected
PA: primary aldosteronism
SP: substance P
ZG: zona glomerulosa
RAS: renin-angiotensin system
BAH: bilateral adrenal hyperplasia

## REFERENCES

1. Monticone S, Burrello J, Tizzani D, et al. Prevalence and Clinical Manifestations of Primary Aldosteronism Encountered in Primary Care Practice. J Am Coll Cardiol. 2017;69(14):1811–1820. doi:10.1016/j.jacc.2017.01.052

2. Buffolo F, Monticone S, Tetti M, Mulatero P. Primary aldosteronism in the primary care setting. Curr Opin Endocrinol Diabetes Obes. 2018;25(3):155–159. doi:10.1097/MED.0000000000000408

3. Funder JW, Carey RM, Mantero F, et al. The Management of Primary Aldosteronism: Case Detection, Diagnosis, and Treatment: An Endocrine Society Clinical Practice Guideline. J Clin Endocrinol Metab. 2016;101(5):1889–1916. doi:10.1210/jc.2015-4061

4. Conn JW. Primary aldosteronism. J Lab Clin Med. 1955;45(4):661–664.

5. Monticone S, D’Ascenzo F, Moretti C, et al. Cardiovascular events and target organ damage in primary aldosteronism compared with essential hypertension: a systematic review and meta-analysis. Lancet Diabetes Endocrinol. 2018;6(1):41–50. doi:10.1016/S2213-8587(17)30319-4

6. Pechère-Bertschi A, Herpin D, Lefebvre H. SFE/SFHTA/AFCE consensus on primary aldosteronism, part 7: Medical treatment of primary aldosteronism. Ann Endocrinol (Paris). 2016;77(3):226–234. doi:10.1016/j.ando.2016.01.010

7. Williams TA, Lenders JWM, Mulatero P, et al. Outcomes after adrenalectomy for unilateral primary aldosteronism: an international consensus on outcome measures and analysis of remission rates in an international cohort. Lancet Diabetes Endocrinol. 2017;5(9):689–699. doi:10.1016/S2213-8587(17)30135-3

8. Williams TA, Gomez-Sanchez CE, Rainey WE, et al. International Histopathology Consensus for Unilateral Primary Aldosteronism. J Clin Endocrinol Metab. 2021;106(1):42–54. doi:10.1210/clinem/dgaa484

9. Fernandes-Rosa FL, Williams TA, Riester A, et al. Genetic spectrum and clinical correlates of somatic mutations in aldosterone-producing adenoma. Hypertension. 2014;64(2):354–361. doi:10.1161/HYPERTENSIONAHA.114.03419

10. Sousa KD, Boulkroun S, Baron S, et al. Genetic, Cellular, and Molecular Heterogeneity in Adrenals With Aldosterone-Producing Adenoma. Hypertension. 2020;75(4):1034. doi:10.1161/HYPERTENSIONAHA.119.14177

11. Choi M, Scholl UI, Yue P, et al. K+ channel mutations in adrenal aldosterone-producing adenomas and hereditary hypertension. Science. 2011;331(6018):768–772. doi:10.1126/science.1198785

12. Beuschlein F, Boulkroun S, Osswald A, et al. Somatic mutations in ATP1A1 and ATP2B3 lead to aldosterone-producing adenomas and secondary hypertension. Nat Genet. 2013;45(4):440–444, 444e1-2. doi:10.1038/ng.2550

13. Azizan EAB, Poulsen H, Tuluc P, et al. Somatic mutations in ATP1A1 and CACNA1D underlie a common subtype of adrenal hypertension. Nat Genet. 2013;45(9):1055–1060. doi:10.1038/ng.2716

14. Scholl UI, Goh G, Stölting G, et al. Somatic and germline CACNA1D calcium channel mutations in aldosterone-producing adenomas and primary aldosteronism. Nat Genet. 2013;45(9):1050–1054. doi:10.1038/ng.2695

15. Scholl UI, Stölting G, Nelson-Williams C, et al. Recurrent gain of function mutation in calcium channel CACNA1H causes early-onset hypertension with primary aldosteronism. Elife. 2015;4:e06315. doi:10.7554/eLife.06315

16. Scholl UI, Stölting G, Schewe J, et al. CLCN2 chloride channel mutations in familial hyperaldosteronism type II. Nat Genet. 2018;50(3):349–354. doi:10.1038/s41588-018-0048-5

17. Fernandes-Rosa FL, Daniil G, Orozco IJ, et al. A gain-of-function mutation in the CLCN2 chloride channel gene causes primary aldosteronism. Nat Genet. 2018;50(3):355–361. doi:10.1038/s41588-018-0053-8

18. van Rooyen D, Bandulik S, Coon G, et al. Somatic Mutations in MCOLN3 in Aldosterone-Producing Adenomas cause Primary Aldosteronism. bioRxiv. October 2024:2024.10.20.619295. doi:10.1101/2024.10.20.619295

19. Berthon A, Drelon C, Ragazzon B, et al. WNT/β-catenin signalling is activated in aldosterone-producing adenomas and controls aldosterone production. Hum Mol Genet. 2014;23(4):889–905. doi:10.1093/hmg/ddt484

20. De Sousa K, Abdellatif AB, Giscos-Douriez I, et al. Colocalization of Wnt/β-Catenin and ACTH Signaling Pathways and Paracrine Regulation in Aldosterone-producing Adenoma. J Clin Endocrinol Metab. 2022;107(2):419–434. doi:10.1210/clinem/dgab707

21. Wu VC, Wang SM, Chueh SCJ, et al. The prevalence of CTNNB1 mutations in primary aldosteronism and consequences for clinical outcomes. Sci Rep. 2017;7:39121. doi:10.1038/srep39121

22. St-Jean M, Bourdeau I, Martin M, Lacroix A. Aldosterone is Aberrantly Regulated by Various Stimuli in a High Proportion of Patients with Primary Aldosteronism. J Clin Endocrinol Metab. 2021;106(1):e45–e60. doi:10.1210/clinem/dgaa703

23. Lefebvre H, Cartier D, Duparc C, et al. Characterization of serotonin(4) receptors in adrenocortical aldosterone-producing adenomas: in vivo and in vitro studies. J Clin Endocrinol Metab. 2002;87(3):1211–1216. doi:10.1210/jcem.87.3.8327

24. Castinetti F, Guerin C, Louiset E, Lacroix A. HCG-responsive aldosteronoma with transient secretion during pregnancy confirmed through HCG-stimulated adrenal venous sampling. Front Endocrinol (Lausanne). 2023;14:1153374. doi:10.3389/fendo.2023.1153374

25. Nichols ML, Allen BJ, Rogers SD, et al. Transmission of chronic nociception by spinal neurons expressing the substance P receptor. Science. 1999;286(5444):1558–1561. doi:10.1126/science.286.5444.1558

26. Saito R, Takano Y, Kamiya HO. Roles of substance P and NK(1) receptor in the brainstem in the development of emesis. J Pharmacol Sci. 2003;91(2):87–94. doi:10.1254/jphs.91.87

27. Prague JK, Roberts RE, Comninos AN, et al. Neurokinin 3 receptor antagonism as a novel treatment for menopausal hot flushes: a phase 2, randomised, double-blind, placebo-controlled trial. Lancet. 2017;389(10081):1809–1820. doi:10.1016/S0140-6736(17)30823-1

28. Steinhoff MS, von Mentzer B, Geppetti P, Pothoulakis C, Bunnett NW. Tachykinins and Their Receptors: Contributions to Physiological Control and the Mechanisms of Disease. Physiological Reviews. 2014;94(1):265–301. doi:10.1152/physrev.00031.2013

29. Wils J, Duparc C, Cailleux AF, et al. The neuropeptide substance P regulates aldosterone secretion in human adrenals. Nature Communications. 2020;11(1):2673. doi:10.1038/s41467-020-16470-8

30. Itcho K, Oki K, Kobuke K, et al. Aberrant G protein-receptor expression is associated with DNA methylation in aldosterone-producing adenoma. Mol Cell Endocrinol. 2018;461:100–104. doi:10.1016/j.mce.2017.08.019

31. Douillard C, Houillier P, Nussberger J, Girerd X. SFE/SFHTA/AFCE Consensus on Primary Aldosteronism, part 2: First diagnostic steps. Ann Endocrinol (Paris). 2016;77(3):192–201. doi:10.1016/j.ando.2016.02.003

32. Lefebvre H, Contesse V, Delarue C, et al. Serotonin-induced stimulation of cortisol secretion from human adrenocortical tissue is mediated through activation of a serotonin4 receptor subtype. Neuroscience. 1992;47(4):999–1007. doi:10.1016/0306-4522(92)90047-6

33. Boyer HG, Wils J, Renouf S, et al. Dysregulation of Aldosterone Secretion in Mast Cell-Deficient Mice. Hypertension. 2017;70(6):1256–1263. doi:10.1161/HYPERTENSIONAHA.117.09746

34. Veldhuis JD, Keenan DM, Pincus SM. Motivations and methods for analyzing pulsatile hormone secretion. Endocr Rev. 2008;29(7):823–864. doi:10.1210/er.2008-0005

35. Silva MSB, Decoster L, Delpouve G, et al. Overactivation of GnRH neurons is sufficient to trigger polycystic ovary syndrome-like traits in female mice. EBioMedicine. 2023;97:104850. doi:10.1016/j.ebiom.2023.104850

36. Shimizu Y, Matsuyama H, Shiina T, Takewaki T, Furness JB. Tachykinins and their functions in the gastrointestinal tract. Cell Mol Life Sci. 2008;65(2):295–311. doi:10.1007/s00018-007-7148-1

37. Heym C, Braun B, Shuyi Y, Klimaschewski L, Colombo-Benkmann M. Immunohistochemical correlation of human adrenal nerve fibres and thoracic dorsal root neurons with special reference to substance P. Histochem Cell Biol. 1995;104(3):233–243.

38. Colombo-Benkmann M, Klimaschewski L, Heym C. Immunohistochemical heterogeneity of nerve cells in the human adrenal gland with special reference to substance P. J Histochem Cytochem. 1996;44(4):369–375.

39. Burnstock G. Autonomic neurotransmission: 60 years since sir Henry Dale. Annu Rev Pharmacol Toxicol. 2009;49:1–30. doi:10.1146/annurev.pharmtox.052808.102215

40. Dehlin HM, Levick SP. Substance P in heart failure: the good and the bad. Int J Cardiol. 2014;170(3):270–277. doi:10.1016/j.ijcard.2013.11.010

41. Hinson JP, Vinson GP, Kapas S, Teja R. The relationship between adrenal vascular events and steroid secretion: the role of mast cells and endothelin. J Steroid Biochem Mol Biol. 1991;40(1-3):381–389.

42. Newby DE, Sciberras DG, Ferro CJ, et al. Substance P-induced vasodilatation is mediated by the neurokinin type 1 receptor but does not contribute to basal vascular tone in man. Br J Clin Pharmacol. 1999;48(3):336–344.

43. Hinson JP, Kapas S, Teja R, Vinson GP. Effect of the endothelins on aldosterone secretion by rat zona glomerulosa cells in vitro. J Steroid Biochem Mol Biol. 1991;40(1-3):437–439. doi:10.1016/0960-0760(91)90213-o

44. Li Q, Johansson H, Kjellman M, Grimelius L. Neuroendocrine differentiation and nerves in human adrenal cortex and cortical lesions. APMIS. 1998;106(8):807–817. doi:10.1111/j.1699-0463.1998.tb00227.x

45. Tischler AS. Divergent differentiation in neuroendocrine tumors of the adrenal gland. Semin Diagn Pathol. 2000;17(2):120–126.

46. Caroccia B, Fassina A, Seccia TM, et al. Isolation of human adrenocortical aldosterone-producing cells by a novel immunomagnetic beads method. Endocrinology. 2010;151(3):1375–1380. doi:10.1210/en.2009-1243

47. Mazzuco TL, Grunenwald S, Lampron A, Bourdeau I, Lacroix A. Aberrant hormone receptors in primary aldosteronism. Horm Metab Res. 2010;42(6):416–423. doi:10.1055/s-0029-1243602

48. Duparc C, Moreau L, Dzib JFG, et al. Mast cell hyperplasia is associated with aldosterone hypersecretion in a subset of aldosterone-producing adenomas. J Clin Endocrinol Metab. 2015;100(4):E550–560. doi:10.1210/jc.2014-3660

49. Campbell DE, Raftery N, Tustin R, et al. Measurement of plasma-derived substance P: biological, methodological, and statistical considerations. Clin Vaccine Immunol. 2006;13(11):1197–1203. doi:10.1128/CVI.00174-06

50. Garcia-Recio S, Gascón P. Biological and Pharmacological Aspects of the NK1-Receptor. Biomed Res Int. 2015;2015:495704. doi:10.1155/2015/495704

51. Page NM, Bell NJ, Gardiner SM, et al. Characterization of the endokinins: Human tachykinins with cardiovascular activity. Proc Natl Acad Sci U S A. 2003;100(10):6245–6250. doi:10.1073/pnas.0931458100

52. Maggi CA, Schwartz TW. The dual nature of the tachykinin NK1 receptor. Trends Pharmacol Sci. 1997;18(10):351–355. doi:10.1016/s0165-6147(97)01107-3

53. Siragy HM, Vieweg WV, Pincus S, Veldhuis JD. Increased disorderliness and amplified basal and pulsatile aldosterone secretion in patients with primary aldosteronism. J Clin Endocrinol Metab. 1995;80(1):28–33. doi:10.1210/jcem.80.1.7829626

54. Oster H, Damerow S, Kiessling S, et al. The circadian rhythm of glucocorticoids is regulated by a gating mechanism residing in the adrenal cortical clock. Cell Metab. 2006;4(2):163–173. doi:10.1016/j.cmet.2006.07.002

55. Oster H, Challet E, Ott V, et al. The Functional and Clinical Significance of the 24-Hour Rhythm of Circulating Glucocorticoids. Endocr Rev. 2017;38(1):3–45. doi:10.1210/er.2015-1080

56. Dibner C, Schibler U, Albrecht U. The mammalian circadian timing system: organization and coordination of central and peripheral clocks. Annu Rev Physiol. 2010;72:517–549. doi:10.1146/annurev-physiol-021909-135821

57. Ehrhart-Bornstein M, Hinson JP, Bornstein SR, Scherbaum WA, Vinson GP. Intraadrenal interactions in the regulation of adrenocortical steroidogenesis. Endocr Rev. 1998;19(2):101–143. doi:10.1210/edrv.19.2.0326

58. Ishida A, Mutoh T, Ueyama T, et al. Light activates the adrenal gland: timing of gene expression and glucocorticoid release. Cell Metab. 2005;2(5):297–307. doi:10.1016/j.cmet.2005.09.009

